# Yeast Bax Inhibitor (Bxi1p/Ybh3p) is a Calcium Channel in *E. coli*

**DOI:** 10.1101/722926

**Authors:** James Mullin, John Kalhorn, Nicholas Mello, Amanda Raffa, Alexander Strakosha, O.P. Nicanor Austriaco

**Affiliations:** Department of Biology, Providence College, Providence, Rhode Island

## Abstract

Human Bax Inhibitor-1 (HsBI-1/TMBIM6) is the founding member of the evolutionary conserved TMBIM superfamily of proteins that share sequence homology within the transmembrane Bax inhibitor-containing motif (TMBIM). Mechanistically, BI-1/TMBIM6 and all the other mammalian TMBIM proteins appear to be involved in the maintenance of calcium homeostasis, and the crystal structure of a bacterial TMBIM protein, BsYetJ, suggests that the protein is a pH-sensitive calcium leak. The budding yeast, *Saccharomyces cerevisiae*, has a single TMBIM family member (YNL305C) named Bxi1p/Ybh3p. To determine the function of Bxi1p/Ybh3p, we overexpressed Bxi1p-EGFP in *E. coli* to determine if it is a calcium channel. We show that bacterial cells expressing Bxi1p-EGFP are more permeable to calcium than controls. Thus, our data suggests that yeast Bax inhibitor (Bxi1p) is a calcium channel in *E. coli*, lending support to our proposal that Bxi1p is a *bona fide* member of the TMBIM family of proteins. Further, we use our bacterial system to show that gadolinium is an inhibitor of Bxi1p *in vivo*, suggesting a path forward to identifying other small-molecular inhibitors of this clinically-important and highly conserved superfamily of proteins. Finally, parallel experiments revealed that the human Bax Inhibitor-1 (HsBI-1/TMBIM6) is also a calcium channel in bacteria that can be inhibited by gadolinium.

## Introduction

Human Bax Inhibitor-1 (HsBI-1/TMBIM6) is the founding member of the evolutionary conserved TMBIM superfamily of proteins that share sequence homology within the transmembrane Bax inhibitor-containing motif (TMBIM) (Xu & Reed, 1998; Reimers et al., 2006; Hu, Smith & Goldberger, 2009; Carrara et al., 2012; Rojas-Rivera & Hetz, 2015; Gamboa-Tuz et al., 2018). The human genome encodes six members of the superfamily (TMBIM1-6) that are homologous to other BI-1 like proteins in vertebrates, plants, yeast, bacteria and viruses. TMBIM1-3 are localized to the Golgi apparatus, TMBIM4-6 are found in the endoplasmic reticulum, and TMBIM5 is a mitochondrial protein (Lisak et al., 2015).

Phenotypically, BI-1/TMBIM6 is an ER-localized, anti-apoptotic protein that was first identified in a screen for human proteins that could inhibit Bax-mediated cell death in yeast (Xu & Reed, 1998). Mammalian cells stably overexpressing BI-1/TMBIM6 are protected against ER-stress induced apoptosis (Chae et al., 2004; Bailly-Maitre et al., 2006). Moreover, mice lacking BI-1/TMBIM6 are more sensitive to stroke-induced cerebral damage and tunicamycin-induced kidney toxicity (Chae et al., 2004). Clinically, BI-1 is known to be overexpressed in breast cancer, glioma, lung, and prostate carcinoma (van’t Veer et al., 2002; Schmits et al., 2002; Grzmil et al., 2003, 2006; Lu et al., 2015). Strikingly, downregulation of BI-1 in prostate cancer cells by RNAi leads to cell death (Grzmil et al., 2003). The other mammalian TMBIM proteins are also cytoprotective against different triggers known to induce cell death (Rojas-Rivera & Hetz, 2015).

Mechanistically, BI-1/TMBIM6 and all the other mammalian TMBIM proteins appear to be involved in the maintenance of calcium homeostasis (Lisak et al., 2015; Liu, 2017). Knocking out TMBIM6 in hepatocytes leads to higher Ca^2+^ content in the ER of hepatocytes (Chae et al., 2004), while overexpressing the gene leads to reduced ER Ca^2+^ content (Westphalen et al., 2005).Similarly, stably overexpressing each HA-tagged TMBIM1-6 protein family member in HT22 cells reduced ER Ca^2+^ content, and all the TMBIM proteins except TMBIM5 also reduced the basal concentration of calcium in the cytosol (Lisak et al., 2015). TMBIM6 regulates Ca^2+^ flux in a pH-dependent manner (Ahn et al., 2009, 2010; Kiviluoto et al., 2013).

Structurally, the role of the TMBIM family of proteins in calcium homeostasis has been confirmed by the solution of the crystal structure of a bacterial TMBIM protein, BsYetJ, from *Bacillus subtilis* that suggests that the protein is a pH-sensitive calcium leak (Chang et al., 2014). It has a seven-transmembrane-helix fold structure that has either a closed or an open channel conformation depending upon the pH of its environment. It also has a di-aspartyl pH sensor in its C-terminal pore domain (Asp171-Asp195) that corresponds to two aspartate residues in BI-1/TMBIM6 (Asp188-Asp213) (Chang et al., 2014). Biochemical characterization of BsYetJ proteoliposomes at various pHs revealed that the pH-sensitive calcium-leak activity is intrinsic to the protein (Chang et al., 2014).

The budding yeast, *Saccharomyces cerevisiae*, has a single TMBIM family member (YNL305C) named Bxi1p/Ybh3p (Chae et al., 2003; Cebulski et al., 2011; Büttner et al., 2011). The protein is homologous to the mammalian TMBIM1-6 family members and contains the conserved di-aspartyl pH sensor in its C-terminal pore domain (Asp255-Asp278) that has been identified as the latch responsible for opening and closing the calcium leak. In a previous study, we showed that Bxi1p-GFP is localized to the ER, and that mutant yeast cells deleted of *BXI1* are more susceptible to a range of pharmacological and environmental triggers that induce cell death, especially pharmacological triggers associated with the unfolded protein response (Cebulski et al., 2011). This pro-survival function for Bxi1p has been confirmed by two other laboratories (Chae et al., 2003; Teng et al., 2011), and is consistent with the anti-apoptotic function associated with the TMBIM superfamily. However, a subsequent paper from the Madeo Laboratory at the University of Graz, published days after our original publication, suggested that the protein encoded by the ORF – which they called Ybh3p for yeast BH3-only protein – is a pro-apoptotic member of the BH3-only family of proteins that translocates from the vacuole to the mitochondria to trigger BH3-domain dependent apoptosis (Büttner et al., 2011).

To begin to experimentally resolve the apparent discrepancy between our data that suggests that Bxi1p/Ybh3p is an anti-apoptotic member of the TMBIM superfamily, and the data that proposed instead that Bxi1p/Ybh3p is a pro-apoptotic member of the BH3-only family of proteins, we overexpressed Bxi1p-GFP in *E. coli* to interrogate its putative calcium channel function. We show that bacterial cells expressing Bxi1p-GFP are more permeable to calcium than controls. Our data suggests that yeast Bax inhibitor (Bxi1p) is a calcium channel in *E. coli*, lending support to our proposal that Bxi1p is a *bona fide* member of the TMBIM family of proteins. We also use our bacterial system to show that gadolinium is an inhibitor of Bxi1p, suggesting a path forward to identifying other small-molecular inhibitors of this clinically-important and highly conserved superfamily of proteins.

## Materials and Methods

### Bacterial Strain, Plasmids, and Growth Conditions

All experiments were done with DH5α *E. coli* cells obtained from New England Biolabs. Bacterial plasmids overexpressing yBxi1p-EGFP and the hTMBIM6-EGFP fusion were constructed and verified by VectorBuilder. The vector ID for plasmid, pBAD-EGFP(ns):4XGS:{yBXI1}, which overexpresses wildtype yBxi1p-EGFP in media containing arabinose is VB160522-1016qch; and the vector ID for plasmid, pBAD-EGFP(ns):4XGS:{hTMBIM6}, which overexpresses hTMBIM6-EGFP is VB170103-1020maq. These vector IDs can be used to retrieve detailed information and plasmid maps from vectorbuilder.com. Cells were transformed and cultured using standard bacterial protocols and media (Ausubel et al., 2002). Unless noted otherwise, all drugs and reagents were purchased from SIGMA-Aldrich.

### Bxi1p-GFP Induction Assays

DH5α *E. coli* cells transformed with a bacterial plasmid overexpressing either yBxi1p-EGFP or hTMBIM6-EGFP were inoculated in 20mL of LB containing 100μg/mL ampicillin and allowed to grow to an OD_600_ of 0.6 at 37°C. Expression of the Bxilp-EGFP fusion protein was then initiated by the addition of 200μL of 20% arabinose. Expression levels of the yBxi1p-EGFP or hTMBIM6-EGFP fusion proteins were then visualized, 1, 3, 6, 12, 18, and 24 hours after induction with a Zeiss LSM700 confocal microscope and quantified with an Accuri C6 Flow Cytometer. For all measurements with the Accuri C6 Flow Cytometer, bacterial cells were first diluted in 10mL water to an OD_600_ of 0.003.

### Calcium Permeability Assays

DH5α *E. coli* cells transformed with a bacterial plasmid overexpressing either yBxi1p-EGFP or hTMBIM6-EGFP were inoculated in 20mL of LB containing 100μg/mL ampicillin and cultured overnight at 37°C in a floor shaker set to 250 RPM. The following day, 1mL of cells were re-inoculated into 10mLs of fresh LB amp and allowed to grow to an OD_600_ of 0.60. Both induced (with 2% arabinose) and uninduced cultures then were incubated at 37°C, 250 RPM for six hours. Next, the cells were spun down at 3,000g for 10mins, washed twice with 10mL Buffer A (50mM Tris pH 7.5; 100mM KCl; 1mM MgCl_2_), and resuspended in 20mL Buffer B (120mM Tris pH 8.0; 0.2mM EDTA) where they were incubated at 37°C, 250 RPM for ten minutes. The cells were then spun down at 3,000g for 10 mins and washed twice with 10mL of buffer A. Finally, the cells were re-suspended in 2.5mL Buffer A containing 10 μM Fura-2AM and incubated at 37°C, 250 RPM for two hours.

Following Fura-2AM incubation, cells were pelleted at 3,000g for 10mins and washed twice with 10mL aliquots of buffer A. Following the washes, the cells were re-suspended in 4mL Buffer A, and plated in 250 μL aliquots in a 96-well black Costar plate. Cells were excited at the standard wavelength for Fura-2AM (510 nm) and measured for wavelength emission of the bound (340nm) and unbound (380 nm) state using a Biotek Cytation 3 Cell Imaging Reader. Emissions were assessed every two minutes for six minutes. Following the initial six minutes, 5 μL of 500 mM CaCl_2_ was added and run at the same conditions for an additional 10 minutes. Calcium concentration was represented by the ratio of R_340_/R_380_ of Fura-2 as has been typically done in previous studies (Hudson et al., 1998; Luo et al., 2019). All experiments were done in triplicate. Statistical significance was determined with the Student’s t-test, using Graph Pad Prism 6. By default, one asterisk is p<0.05; two asterisks is p<0.01; three asterisks is p<0.001; and four asterisks is p<0.0001.

### Gadolinium Assays

To determine whether or not gadolinium inhibited calcium channel activity, the calcium permeability assay described above was repeated with an additional step: Following the Fura-2AM incubation and the emission baseline measurements, but before the addition of external calcium, 10μL of 500 mM GdCl_3_ was added to the cells in the 96-well plates for 10 minutes to determine if the larger cation had any effect on the cell’s permeability to Ca^2+^.

To determine whether or not gadolinium had any effect on bacterial cell growth, DH5α *E. coli* cells transformed with the appropriate vector were streaked onto LB-ampicillin plates with varying concentrations of gadolinium (GdCl_3_). The plates were incubated at 37°C for two days and imaged with a SynGene G:BOX system.

## Results

### Yeast Bax Inhibitor (Bxi1p/Ybh3p) is a Calcium Channel in *E. coli*

To determine if yeast Bax inhibitor (Bxi1p) has calcium channel function in *E. coli*, we overexpressed a Bxi1p-EGFP fusion protein in bacterial cells using the arabinose-inducible araBAD promoter (Guzman et al., 1995). A similar experiment had been done with the *Bacillus subtilis* homolog, BsYetJ, in bacteria (Chang et al., 2014).

Expression levels of the Bxi1p-EGFP fusion protein were visualized 1, 3, 6, 12, 18, and 24 hours after induction on a Zeiss LSM700 confocal microscope and were quantified with an Accuri C6 Flow Cytometer to measure GFP fluorescence. As shown in Figures 1A and 1B, bacterial cells grown in media containing arabinose for six hours showed significant induction (p<0.0001) of the Bxi1p-EGFP fusion protein. This time point was chosen for all further experiments. We then loaded these DH5α cells with Fura-2AM, a membrane-permeable, fluorescent calcium indicator that has been used to determine the intracellular calcium dynamics of bacterial cells (Gangola & Rosen, 1987; Chang et al., 2014). Upon addition of external calcium, intracellular calcium concentration as indicated by the R_340_/R_380_ ratio of Fura-2 increased more rapidly in bacterial cells induced with arabinose as compared with uninduced controls (Figure 1C). This data suggests that Bxi1p-EGFP is a functional TMBIM superfamily member with channel activity that can increase the permeability of bacterial cells to calcium.

**FIGURE 1:**
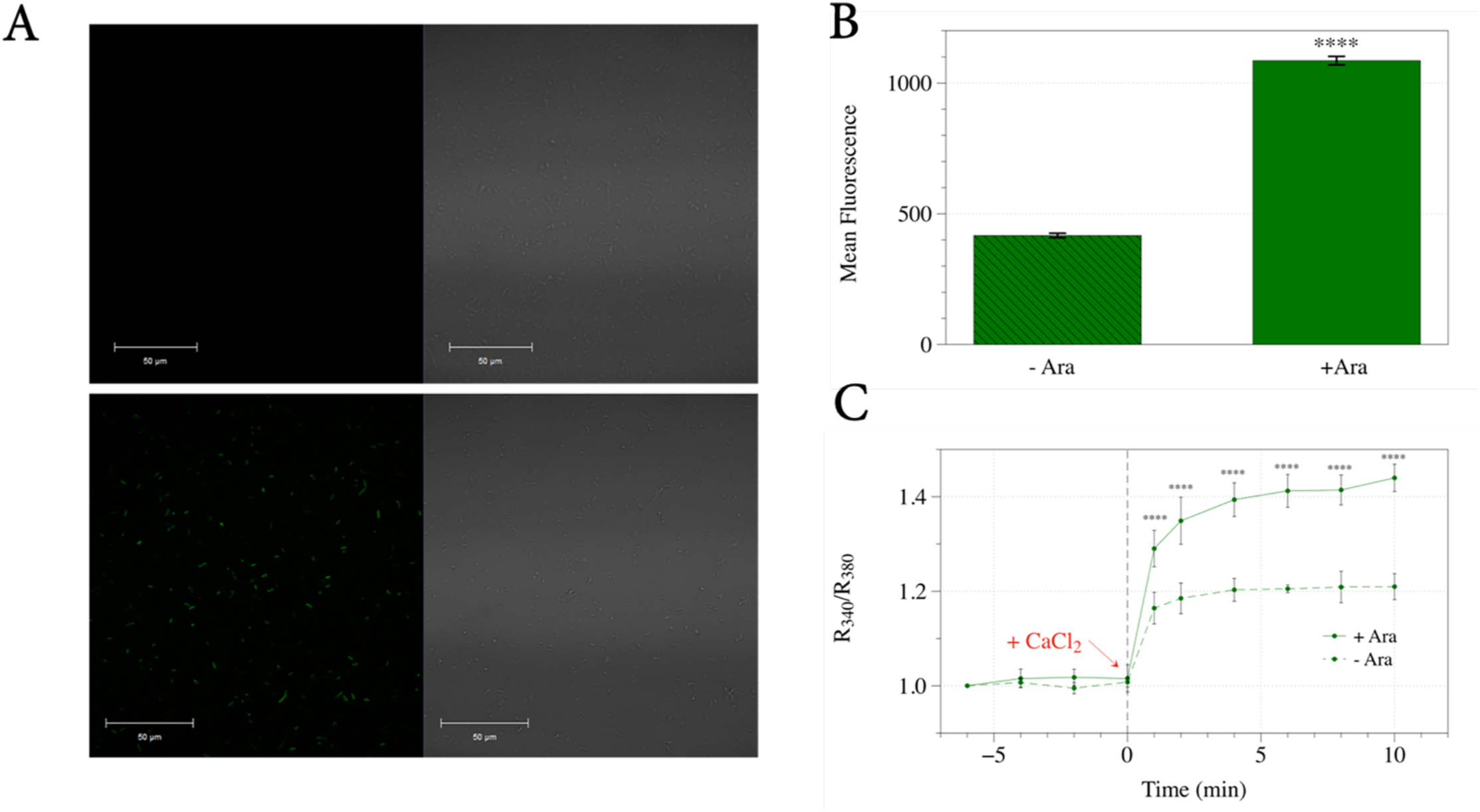
Yeast Bax Inhibitor (Bxi1p/Ybh3p) is a Calcium Channel in *E. coli*. (A and B) We expressed a yeast Bax inhibitor-EGFP fusion protein in DH5α bacterial cells using the arabinose-inducible araBAD promoter. Bacterial cells grown in LB media containing arabinose for six hours showed significant induction of the Bxi1p-EGFP fusion protein as measured by an Accuri C6 Flow Cytometer. (C) We then loaded these DH5α cells with Fura-2AM, a membrane-permeable, fluorescent calcium indicator that has been used to determine the intracellular calcium dynamics of bacterial cells. Upon addition of external calcium, intracellular calcium concentration as indicated by the R340/R380 ratio of Fura-2 increased more rapidly in bacterial cells induced with arabinose as compared with uninduced controls. Error bars indicate standard deviations for trials with at least three independent cultures. The differences in expression and calcium levels were deemed statistically significant by the Student’s t-test (* p<0.05; ** p<0.01; *** p<0.001; **** p<0.0001).

### Yeast Bax Inhibitor’s Channel Function in *E.coli* Can Be Inhibited by Gadolinium

To see if we could identify small-molecule inhibitors of the TMBIM superfamily of calcium channels using our *in vivo* bacterial system overexpressing Bxi1p-EGFP, we tested several ions for their ability to block calcium permeability in our cells, reasoning that cations larger than calcium may be able to inhibit Ca^2+^ binding to Bxi1p. To do this, we repeated our overexpression experiments with an additional incubation step: Before the addition of external calcium, the larger cations were added to wells containing Fura-2AM loaded bacterial cells to see if they had any effect on the cell’s permeability to Ca^2+^.

Of several cations tested, we discovered that the gadolinium ion, Gd^3+^, which is a potent blocker of calcium channels (Bourne & Trifaró, 1982; Biagi & Enyeart, 1990; Malasics et al., 2010) was able to block the permeability of calcium in our DH5α cells with both basal and overexpressed levels of Bxi1p-EGFP (Figure 2A). Cells pre-exposed to Gd^3+^ did not manifest the calcium spike as measured by the R_340_/R_380_ ratio of Fura-2 that was observed in non-treated cells. Importantly, the trivalent cation did not significantly decrease the proteins levels of Bxi1p-EGFP as determined by flow cytometry (Figure 2B), nor did it lead either to slow growth or to bacterial cell death (Figure 2C), suggesting that it is a *bona fide* inhibitor of Bxi1p-EGFP’s function.

**FIGURE 2:**
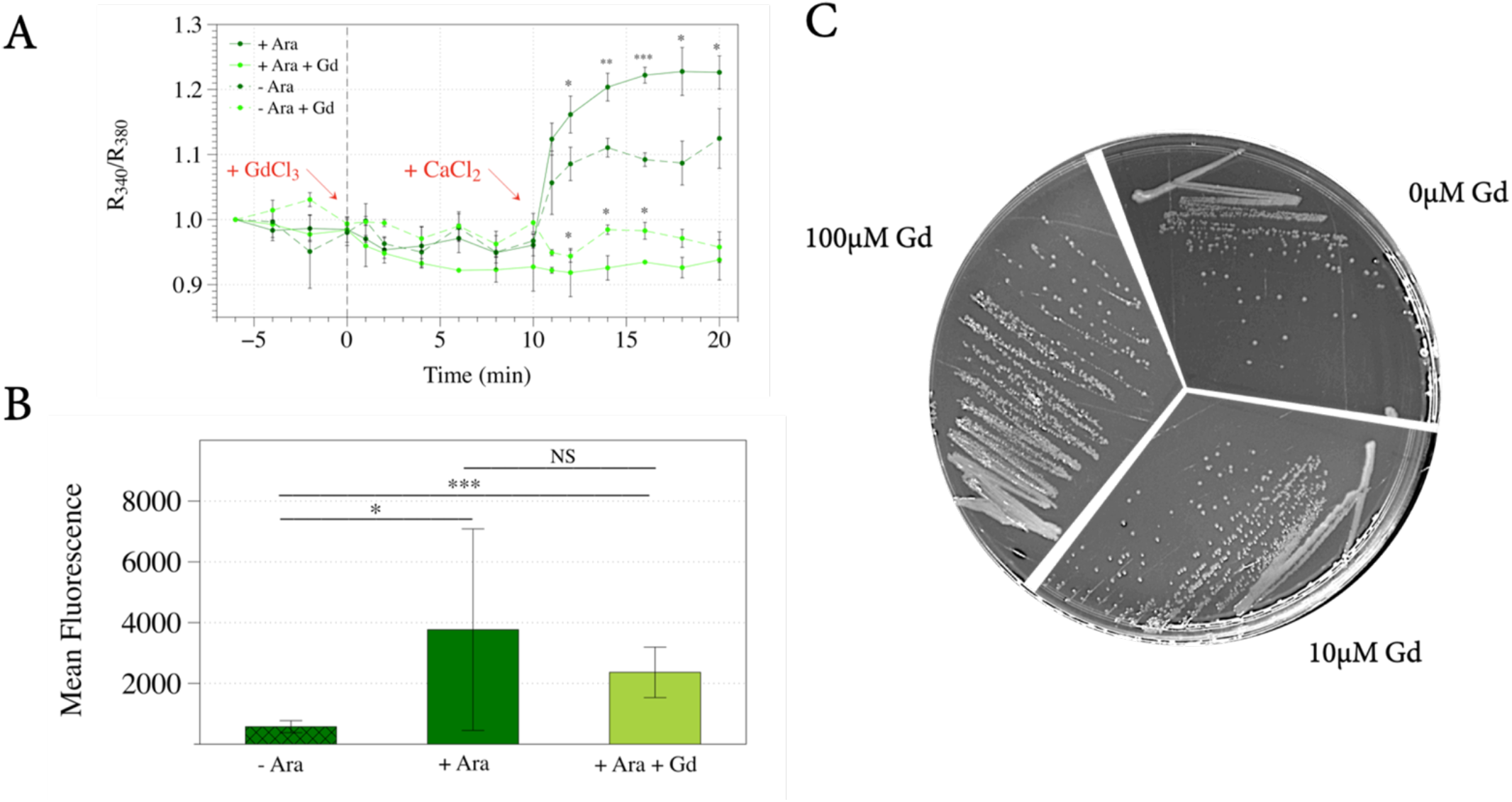
Yeast Bax Inhibitor’s Channel Function in *E.coli* Can Be Inhibited by Gadolinium. (A) Bacterial cells overexpressing the yeast Bax inhibitor, Bxi1p-EGFP fusion protein were grown in LB media with and without arabinose for six hours. Following the six-minute emission baseline measurements, the cells were loaded with 10μL of 500 mM GdCl_3_. Upon addition of external calcium ten minutes later, the increase in intracellular calcium concentration as indicated by the R340/R380 ratio of Fura-2 was inhibited by Gd^3+^ treatment. Error bars indicate standard deviations for trials with at least three independent cultures. The differences in expression levels were deemed statistically significant by the Student’s t-test (* p<0.05; ** p<0.01; *** p<0.001; **** p<0.0001). (B) To determine if gadolinium inhibited expression of the Bxi1p-EGFP fusion, levels of EGFP in bacterial cells grown in arabinose with and without gadolinium were compared with an Accuri C6 Flow Cytometer. Error bars indicate standard deviations for trials with at least three independent cultures. The differences in expression levels were deemed statistically significant by the Student’s t-test (* p<0.05; ** p<0.01; *** p<0.001; **** p<0.0001). (C) To determine whether or not gadolinium had any effect on bacterial cell growth, DH5α *E. coli* cells transformed with the Bxi1p-EGFP expression vector were streaked onto LB-Ampicillin plates with varying concentrations of gadolinium (GdCl_3_). The plates were incubated at 37°C for two days and imaged.

### Human Bax Inhibitor’s Channel Function in *E. coli* Can Also Be Inhibited by Gadolinium

To determine if the human homolog of Bax inhibitor (HsBI-1/TMBIM6) could function as a calcium channel in our bacterial *in vivo* assay, we overexpressed a protein HsBI1-EGFP fusion using the same arabinose-inducible araBAD promoter as was used with the yeast Bxi1p-EGFP homolog described above. As shown in Figure 3A, bacterial cells grown in media containing arabinose for six hours showed significant induction(p<0.05) of the HsBI1-EGFP fusion protein. Upon addition of external calcium, intracellular calcium concentration as indicated by the R_340_/R_380_ ratio of Fura-2 increased more rapidly in cells induced with arabinose as compared with uninduced controls (Figure 3B). This data suggests that HsBI1-EGFP is also a calcium channel when expressed in *E. coli*. This is not unexpected given the significant sequence homology between human Bax inhibitor (hBI-1/TMBIM6) and its bacterial homolog, BsYetJ (Chang et al., 2014).

**FIGURE 3:**
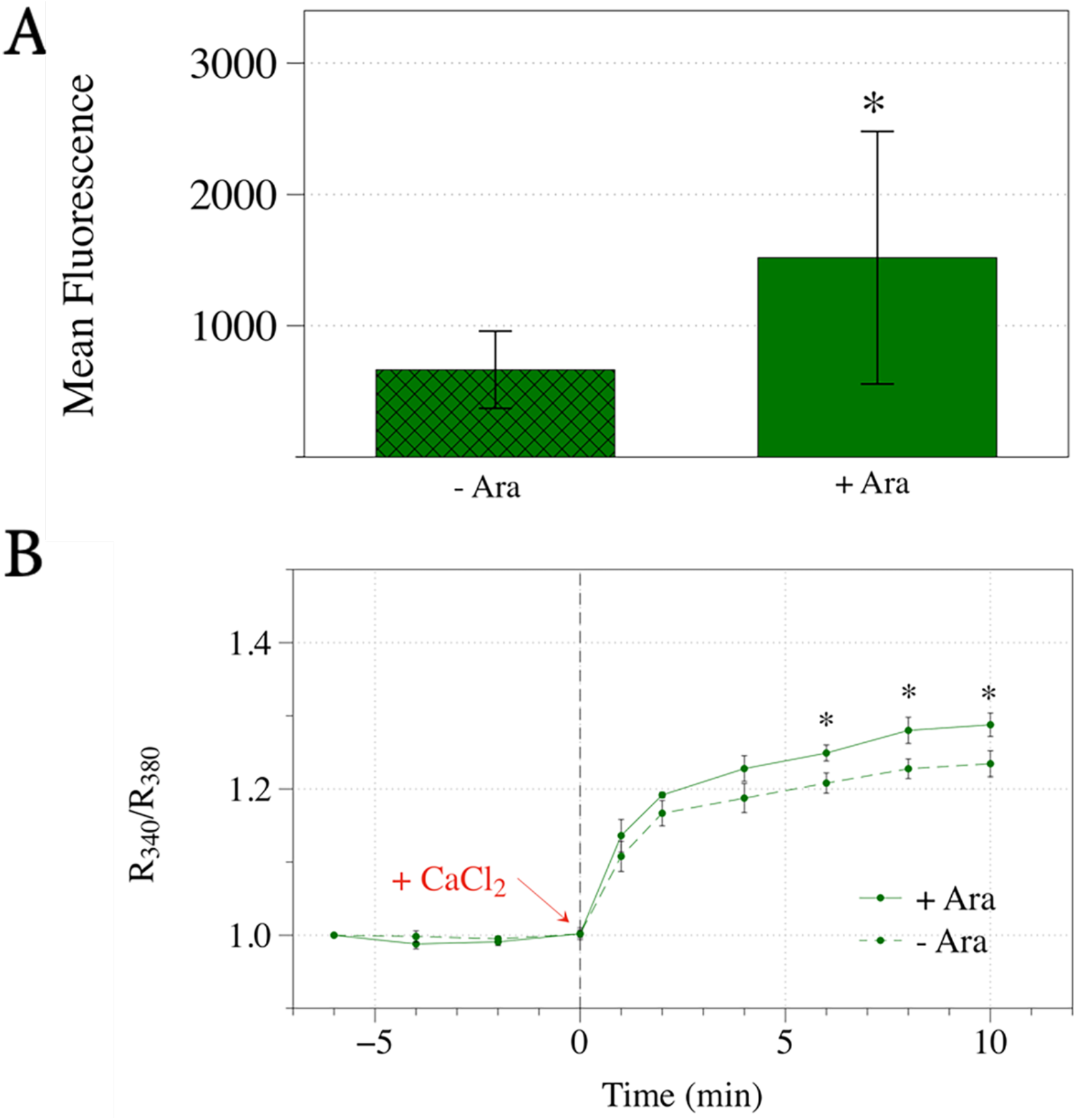
Human Bax Inhibitor (BI-1/TMBIM6) is a Calcium Channel in *E. coli*. (A and B)We overexpressed a human Bax inhibitor, HsBI1-EGFP fusion protein in DH5α bacterial cells using the arabinose-inducible araBAD promoter. Bacterial cells grown in LB media containing arabinose for six hours showed significant induction of the BI1-EGFP fusion protein as measured by an Accuri C6 Flow Cytometer. (C) We then loaded these DH5α cells with Fura-2AM, a membrane-permeable, fluorescent calcium indicator that has been used to determine the intracellular calcium dynamics of bacterial cells. Upon addition of external calcium, intracellular calcium concentration as indicated by the R340/R380 ratio of Fura-2 increased more rapidly in bacterial cells induced with arabinose as compared with uninduced controls. Error bars indicate standard deviations for trials with at least three independent cultures. The differences in expression and calcium levels were deemed statistically significant by the Student’s t-test (* p<0.05; ** p<0.01; *** p<0.001; **** p<0.0001).

Finally, we discovered that Gd^3+^ is also able to block the permeability of calcium in bacterial cells with both basal and overexpressed levels of HsBI1-EGFP (Figure 4A). Once again, the trivalent cation did not decrease the levels of Bxi1p-EGFP as measured by flow cytometry (Figure 4B), nor did it lead either to slow growth or to bacterial cell death (Figure 4C), suggesting that it is a *bona fide* inhibitor of HsBI1-EGFP’s function.

**FIGURE 4:**
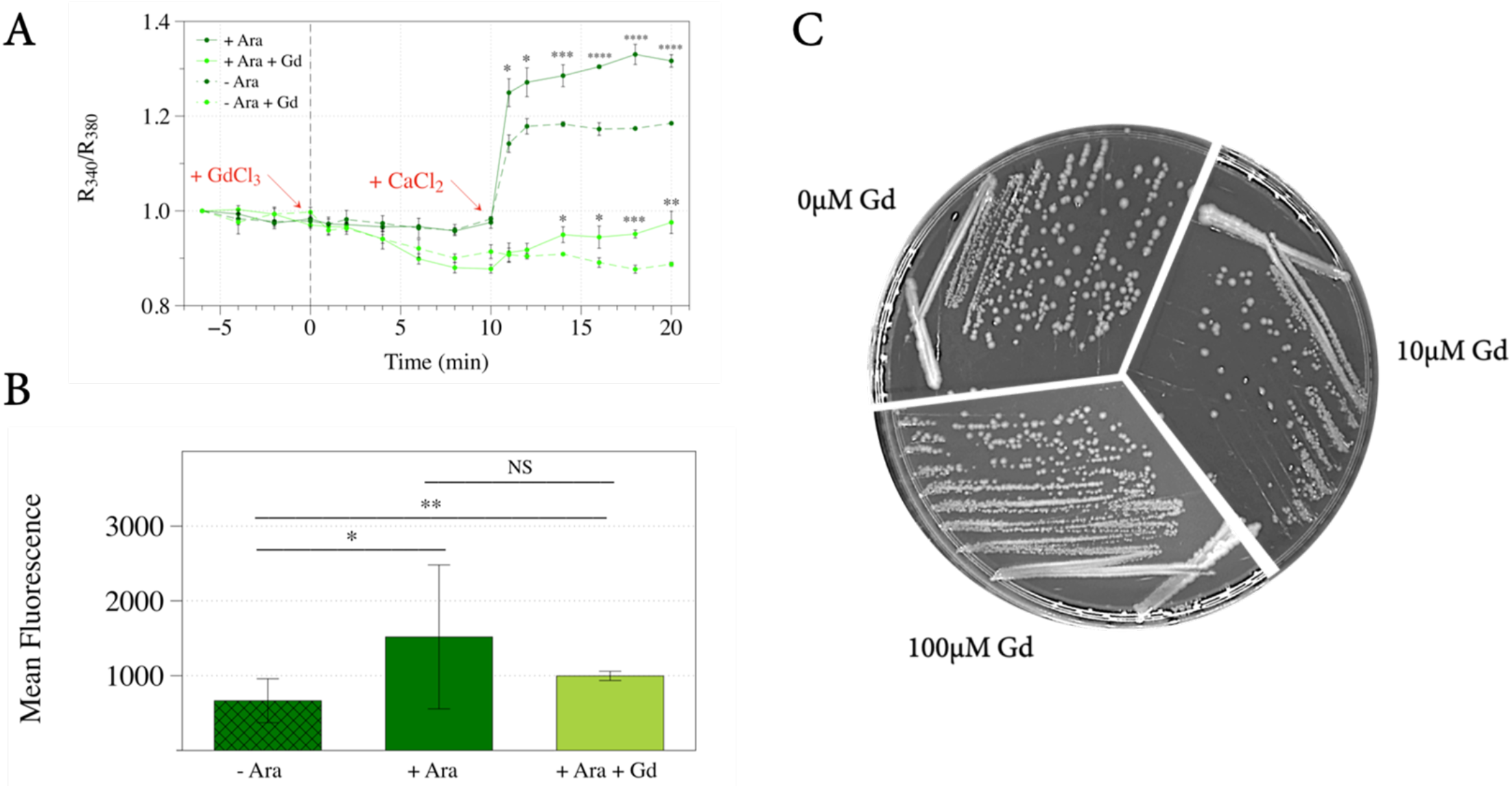
Human Bax Inhibitor’s Channel Function in *E.coli* Can Be Inhibited by Gadolinium. (A) Bacterial cells overexpressing the human Bax inhibitor, HsBI1-EGFP fusion protein were grown in LB media with and without arabinose for six hours. Following the six-minute emission baseline measurements, the cells were loaded with 10μL of 500 mM GdCl_3_. Upon addition of external calcium ten minutes later, the increase in intracellular calcium concentration as indicated by the R340/R380 ratio of Fura-2 was inhibited by Gd^3+^ treatment. Error bars indicate standard deviations for trials with at least three independent cultures. The differences in calcium levels were deemed statistically significant by the Student’s t-test (* p<0.05; ** p<0.01; *** p<0.001; **** p<0.0001). (B) To determine if gadolinium inhibited expression of the HsBI1-EGFP fusion, levels of EGFP in bacterial cells grown in arabinose with and without gadolinium were compared with an Accuri C6 Flow Cytometer. Error bars indicate standard deviations for trials with at least three independent cultures. The differences in expression levels were deemed statistically significant by the Student’s t-test (* p<0.05; ** p<0.01; *** p<0.001; **** p<0.0001). (C) To determine whether or not gadolinium had any effect on bacterial cell growth, DH5α E. coli cells transformed with the HsBI1-EGFP expression vector were streaked onto LB-Ampicillin plates with varying concentrations of gadolinium (GdCl_3_). The plates were incubated at 37°C for two days and imaged.

## Discussion

The budding yeast, *Saccharomyces cerevisiae*, has a single TMBIM family member (YNL305C) named Bxi1p/Ybh3p (Chae et al., 2003; Cebulski et al., 2011; Büttner et al., 2011). However, the precise function of this protein is disputed. In a previous paper, we showed that Bxi1p-GFP is localized to the ER, and that mutant yeast cells deleted of *BXI1* are more susceptible to a range of pharmacological and environmental triggers that induce cell death, especially pharmacological triggers associated with the unfolded protein response (Cebulski et al., 2011). This pro-survival function for Bxi1p has been confirmed by two other laboratories (Chae et al., 2003; Teng et al., 2011), and is consistent with the anti-apoptotic function associated with the TMBIM superfamily.

In contrast, another publication has suggested that the protein encoded by YNL305C – which was named Ybh3p for yeast BH3-only protein – is a pro-apoptotic member of the BH3-only family of proteins that is able to bind *in vitro* with Bcl-X_L_ (Büttner et al., 2011). Moreover, the authors of this paper propose that Ybh3p translocates from the vacuole to the mitochondria to trigger BH3-domain dependent apoptosis. This would suggest that Bxi1p/Ybh3p has a pro-apoptotic function consistent with its proposed membership in the BH3-only family of proteins.

How do we resolve this discrepancy in the reported putative functions of Bxi1p/Ybh3p? It is notable that the inclusion of Bxi1p/Ybh3p in the BH3-only family of proteins has been criticized by those who have proposed that the BH3-domain-like sequence in this yeast ORF is not a *bona fide* BH3 domain (Aouacheria et al., 2013). These critics point out that the candidate yeast BH3 sequence in YNL305C, which was initially identified by visual inspection, is somewhat truncated as it is located at the C-terminus of the Ybh3p protein, and it overlaps one of six transmembrane segments, two unprecedented features among any other known BH3-containing proteins. They also affirm that in principle, it is perplexing for budding yeast to have a BH-3 only protein, since it is not clear how this protein would have ever encountered human Bcl-X_L_ in the course of evolutionary history. Nonetheless, they concede that it is possible that the yeast genome may include a not-yet identified protein whose structure so closely resembles the 3D structure of the Bcl-2 family proteins that it could serve as a potential target of this yeast BH3 domain.

In this paper, we report that the overexpression of either Bxi1p/Ybh3p-EGFP and HsBI1-EGFP increases the permeability of bacterial cells to external calcium. This suggests that Bxi1p-EGFP, like its human counterpart, is a functional TMBIM superfamily member with calcium channel activity. It also suggests that Bxi1p-EGFP is not a member of the BH3-only family of proteins, which are known to be globular rather than transmembrane proteins (Glab, Mbogo & Puthalakath, 2017). However, we are still not sure how to reconcile this observation that Bxi1p is a transmembrane channel protein with the published data that suggests that Bxi1p/Ybh3p is a protein that translocates from the vacuole to the mitochondria to trigger cell death in a manner reminiscent of the BH3-only family of proteins (Büttner et al., 2011).

In addition, we show that the gadolinium ion, Gd^3+^, appears to inhibit both Bxi1p-EGFP and HsBI1-EGFP function in *E. coli*. To the best of our knowledge, this is the first known small-molecule inhibitor of a TMBIM superfamily member that works *in vivo*. However, it is not clear if Gd^3+^ blocks the channel’s pore by binding to the Ca^2+^ binding site or if it alters the conformation of the protein itself to prevent Ca^2+^ binding and channel function.

Notably, as we were about to submit this manuscript for publication, a paper was published that showed that gadolinium inhibits the binding of ^45^Ca^2+^ to the bacterial TMBIM homolog, BsYetJ, *in vitro* (Guo et al., 2019). This report also revealed that Yb^3+^ and Lu^3+^ were able to inhibit Ca^2+^ to the BsYetJ channel, albeit to a much lesser extent. As the authors acknowledged, however, it is still not clear if this inhibition of channel function is the result of direct binding of the lanthanides to the Ca^2+^ binding site or of an induced conformational change in the channel’s tertiary structure triggered by the larger cations.

Finally, given the observation that RNA interference of Bax inhibitor activity in the prostate cancer cell lines, PC-3, LNCaP, and DU-145, triggered cell death (Grzmil et al., 2003), it would be interesting to see if Gd^3+^ treatment of these cell lines would also trigger apoptosis by inhibiting the channel function of the protein. In principle, this would suggest that small molecule inhibitors of TMBIM function could be therapeutic in nature.

## Conclusion

In this paper, we report that the overexpression of either Bxi1p/Ybh3p-EGFP and HsBI1-EGFP increases the permeability of bacterial cells to external calcium and that this activity can be inhibited by gadolinium *in vivo*. It suggests that Bxi1p like its human counterpart is a *bona fide* member of the TMBIM protein family of calcium channels. We are currently using our *in vivo* system to identify other small-molecular inhibitors of this clinically-important and highly conserved superfamily of proteins.

